# Bovine endometrial organoids: A new tool to study conceptus-maternal interactions in mammals

**DOI:** 10.1101/2024.12.13.628419

**Authors:** Jessica C. Edge, Olga Amelkina, Haidee Tinning, Gianluca Giovanardi, Elena Mancinelli, Samantha Gardner, Elton JR Vasconcelos, Virginia Pensabene, Karen Forbes, Mary J O’Connell, Peter Ruane, Niamh Forde

## Abstract

In cattle, communication between endometrium and conceptus during the peri-implantation period is crucial for successful pregnancy. Understanding these interactions is vital as most early pregnancy loss occurs during this time. A major challenge in understanding uterine function and early pregnancy is lack of appropriate *in-vitro* models. Two-dimensional models are available, but do not recapitulate the endometrium’s complex multicellular structure. Here, we describe a hormonally responsive organoid model of the bovine endometrium, developed as a tool for studying endometrial function and early pregnancy. Bovine glandular epithelial cells were isolated from reproductive tracts and cultured in an extracellular matrix hydrogel (Cultrex 2) at 38.5°C, 5% CO_2_ (n=3). RNA was extracted and qPCR confirmed the presence of gland markers: *leukemia inhibitory factor, mucin-1, insulin-like growth factor binding protein-1, kruppel-like factor-5* and *forkhead box protein-A2*. Organoids were imaged at specific time-points to monitor growth and passaged 3 times in 1:2 or 1:3 ratios after growing for a minimum of 10 days per passage. Morphologically, organoids were spherical and fast-growing at passages 0 and 1, but this declines following passage 2. Bovine endometrial organoids (n=3, passage 0) were treated with 1000 ng/ml recombinant ovine Interferon Tau (IFNT) or 10 μg/ml progesterone (P4) for 24 hours and analysed by RNASeq to assess hormone responsiveness. Differential expression analysis by DeSeq2 negative binomial distribution model followed by Wald test and Benjamini-Hochberg correction identified 373 transcripts significantly upregulated (padj<0.05 or log_2_fold change >0.05) in response to P4 treatment, with downstream analysis showing significant overrepresentation (FDR<0.05) of genes associated with positive regulation of protein localisation to plasma membrane and cell periphery. Of the 240 genes significantly downregulated by P4 these were significantly overrepresented (FDR<0.05) in biological processes of cilium and cytoskeleton organisation. IFNT treatment resulted in significant upregulation of 414 genes and downregulation of 119 genes. The largest cluster associated with differentially expressed genes in response to IFNT is defence to virus and interferon signalling. There were 30 genes altered by both P4 treatment and IFNT treatment. Organoids were also shown to express conserved microRNAs, and it was possible to culture them in a microfluidics device - making them a useful model for a multitude of potential investigations. This model provides a tool to investigate bovine endometrial function and peri-implantation communication, subsequently allowing species comparison to understand diversity in reproductive strategies.

## INTRODUCTION

Understanding endometrial function, particularly in domestic species, has been limited by the lack of in vitro models that mimic the in vivo scenario, as well as the cost of in vivo approaches. Whilst in vivo approaches have been used to understand uterine luminal fluid (ULF) composition ^1–3^, actions of critical pregnancy cues like progesterone (P4) and Interferon Tau (IFNT)^3–5^, as well as the biosensor capability of the endometrium^6,7^, it has been difficult to study endometrial function in a high throughout manner. An endometrium that functions correctly is critical to pregnancy success in all mammals, given it is the site of implantation, and it provides nutrient support via ULF to the developing conceptus (embryo and extra-embryonic membranes) prior to development of the placenta. Moreover, dysfunction of the endometrium leads to pregnancy loss. In cattle, embryonic loss most commonly occurs in the first 3 weeks of pregnancy and is detrimental both financially - to the meat and dairy industries; as well as having a negative effect on sustainability and the environment^8^. For these reasons, unravelling how the complex roles of the endometrium in embryo development and conceptus-maternal interactions interplay for successful early pregnancy is imperative.

For successful pregnancy in cattle, an adequately developed, elongated conceptus must meet an optimally primed endometrium^9^. The bovine endometrium is a complex tissue whose transcriptional and translational profile changes throughout the cycle and pregnancy. It comprises a stromal cell layer, with an overlying epithelial cell layer containing uterine glands (both superficial and deep glandular in nature with different phenotypes) which extend into the underlying stromal layer^10^. Factors secreted from the luminal and glandular epithelium are integral for conceptus elongation^11^ highlighting the necessity to understand the components of these secretions.

Until recently, 2D in vitro cultures were the most used approach to study conceptus-maternal interactions^12^ during the peri-implantation period of pregnancy. Many 2D cell culture models utilise cell lines, which can introduce drawbacks. One of the main concerns with using cell lines is that genetic drift or contamination may occur following repeated passaging, potentially altering the physiological behaviour of the cells ^13^. The “normal” functioning of the cell may also be affected due to the fact that cells are often immortalized; or cancer derived cells which behave differently to a non-cancerous cell^14^. Although primary cells can be used to mitigate some of these issues, 2D single cell type monolayers do not accurately recapitulate tissues and organs, introducing the need for more complex in vitro systems. Despite their limitations, these methods are robust and convenient; and if taken tentatively with knowledge of their limitations, provide effective data. For example, treatment with molecules important for early pregnancy can reveal cell type specific functional effects^15^. However, it is undeniable that single cell type, static, in vitro cultures do not represent the complex multicellular endometrium in vivo. A study investigated the impact of using 2D vs 3D culture for bovine uterine gland fragments^16^. The mRNA expression of secretory proteins thought to be involved in conceptus development was significantly lower in 2D culture compared to 3D. Cell morphology in the 3D culture was also more physiologically accurate, with cells exhibiting cubic shapes and displaying apical and basolateral localisation of Zona Occluden 1 (ZO-1) and Beta catenin; whereas 2D culture cells were flat with non-specific localisation. This demonstrates the need to develop 3D endometrial culture approaches that more closely mimic the in vivo situation.

One such approach is the use of in vitro organoids which are self-organising, 3D structures, derived from primary tissue to represent the more complex organ that can be cultured in vitro^17^. Previously used to model various tissues involving epithelial cells including colon^18^, prostate^19^ and fallopian tube^20^; a breakthrough paper developed human endometrial organoids^21^ using both endometrial gland tissue and decidual cells. These endometrial organoids are responsive to key cycle-related hormones P4 and estrogen (E2), can withstand long-term culture and passaging, making them an excellent candidate tool to investigate endometrial function. Collection of these cells requires an invasive endometrial biopsy procedure, however more recently, menstrual flow derived organoids have been produced which respond appropriately to hormonal cues^22^. This advancement improves accessibility and availability in menstruating species. Development of oviductal organoids from other species has also been achieved in cow, pig, dog, and horse^23^. Applying this technology to the endometrium to develop bovine endometrial organoids would provide an additional tool for understanding complex conceptus-maternal interactions in the peri-implantation period of pregnancy. Therefore, the aim of this study was to develop and characterise a method to generate bovine endometrial organoids that respond to key cues important for pregnancy success to facilitate higher throughput studies of endometrial function in vitro.

## MATERIALS AND METHODS

Unless otherwise stated, all reagents were sourced from Sigma (UK). All reproductive tracts were sourced from a local abattoir.

### Optimization of organoid isolation process

Reproductive tracts were selected based on CL morphology as previously described (mid-late-luteal phase^24^). Tracts were washed with 70% ethanol, transferred to a hood and further washed with 70% ethanol (Figure 1A). The ipsilateral horn was identified, and the horn was cut above the uterine body. A cut was made down the horn toward the oviduct to expose the uterine endometrium. The endometrium was washed with 25 ml of endometrial wash buffer (Dulbecco’s Phosphate Buffered Saline (DPBS) and 1% ABAM) dissected from the underlying myometrium and placed in 25 mL of endometrial strip wash (HBSS with no calcium and magnesium and 1% antibiotic, antimycotic (ABAM)). The strip wash was removed and replaced with 25 ml HBSS to wash, before fragments were transferred to a petri dish for further dissection with a scalpel into approximately 1 mm fragments and returned to HBSS before digestion. Three different compositions of digestion solution were used to optimise yield of glandular epithelial cells, differing in concentration of dispase and DNase I. Each digestion composition contained 0.5 mg/ml collagenase II, along with: 1) 100 mg/ml DNase I; 2) 100 mg/ml DNase I and 1 mg/ml dispase; or 3) 1 mg/ml DNase I and 1 mg/ml dispase.

**Figure 1.**
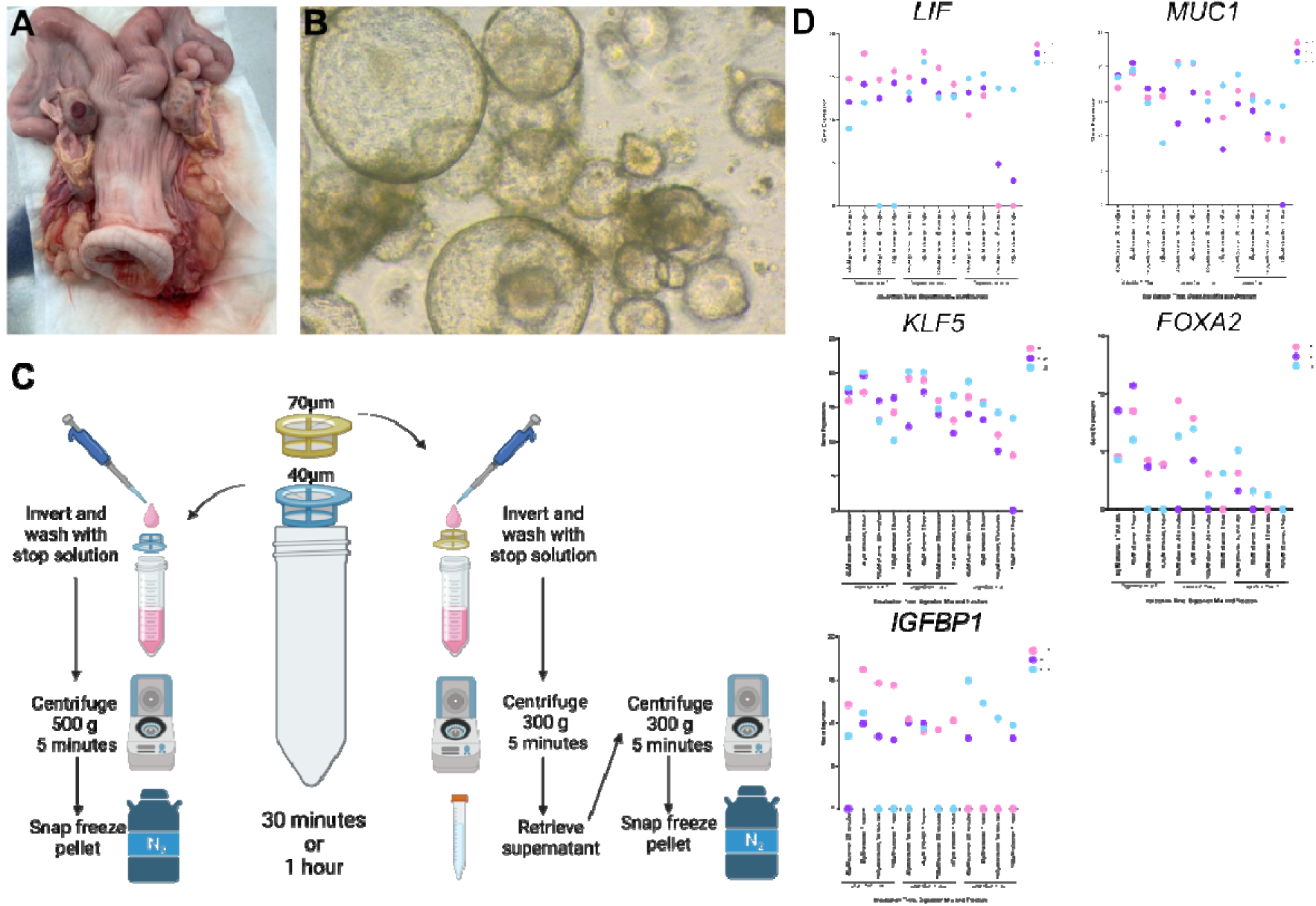
Isolation of bovine endometrial organoids. **(A)** Bovine uterus note the corpus luteum in the mid-secretory phase. **(B)** Light microscopy image of organoids resulting from isolation by digestion method 1 and incubation time of 1 hr. **(C)** Schematic diagram demonstrating workflow of isolating cells to generate bovine endometrial organoids. Cells were isolated using different combinations of digestion solutions and fractions of cell strainers for varying times to determine optimal isolation methods for glandular organoid isolation. **(D)** Comparison of gland markers LIF, MUC1, KLF5, FOXA2, and IGFBP1, in cells using different isolation methods. Gene expression in cells isolated from the 40μM strainer or the 70μM strainer from bovine endometrium (n=3) following either 30 min or 1 hr incubation at 38.5 °C 5% CO2 in digestion mix 1 (0.5mg/ml collagenase II, 100μg/ml DNase), 2 (0.5mg/ml collagenase II, 1mg/ml dispase, 100μg/ml DNase) or 3 (0.5mg/ml collagenase II, 1mg/ml dispase, 1μg/ml DNase).

Each digestion composition was separately subjected to a 30 min and 1 hr incubation at 38.5°C whilst being agitated to optimise length of digestion. After incubation, solutions were filtered through two filters, firstly 100 μM and then 40 μM. Different fractions of the digestion were collected in order to select the one that was most glandular cell enriched. The first fraction was collected by inverting the 40 μM filter and rinsing with 5 ml stop solution (50 ml HBSS and 5 ml Foetal Bovine Serum (FBS)), followed by a 5 min centrifugation at 500 g. Supernatant was aspirated and pellet snap frozen in liquid nitrogen for subsequent analysis. The 100 μM filter was also inverted and rinsed with 5 ml stop solution; this fraction containing tissue chunks was centrifuged for 5 min at 300 g. The supernatant was transferred to another tube, centrifuged for 5 min at 500 g, supernatant aspirated and pellet snap frozen in liquid nitrogen. This process is represented in Figure 1C.

### RNA extraction, cDNA conversion and qRT-PCR analysis

Total RNA was extracted from each fraction, at each time point, using the RNeasy Mini Kit and on-column DNase digestion as per manufacturer’s protocol (Qiagen). RNA concentration was determined using the DeNovix DS-11 FX+ spectrophotometer (DeNovix). Extracted RNA was reverse transcribed using the High-Capacity cDNA reverse transcription kit (Applied Biosystems) as per manufacturer’s protocol. Primers for selected glandular markers were designed using Primer Blast to span an exon-exon junction with a minimum product size of 50 and maximum of 100. Primers with the fewest unintended targets were selected and are listed in Table 1. Quantitative-Realtime PCR (qPCR) was carried out using SYBR green (Roche) with 5 ng cDNA per reaction. qPCR was performed using the LightCycler 96 (Roche) under the following cycling conditions: preincubation at 95 °C for 5 min; 45 cycles of 95 °C for 10 sec, 56 °C for 10 sec, 72 °C for 10 sec; then melting at 95 °C for 5 sec, 65 °C for 1 min and 97 °C continuously. Absolute quantification of data was obtained to collect Ct values using LightCycler96 software before exportation to Microsoft Excel for further analysis. qPCR was carried out in technical triplicate, and the average Ct value of these three measurements was subtracted from 45 (the maximum possible Ct value) to calculate raw gene expression values for the selected markers of endometrial glands. Graphs were plotted using Graphpad Prism.

**Table 1.**
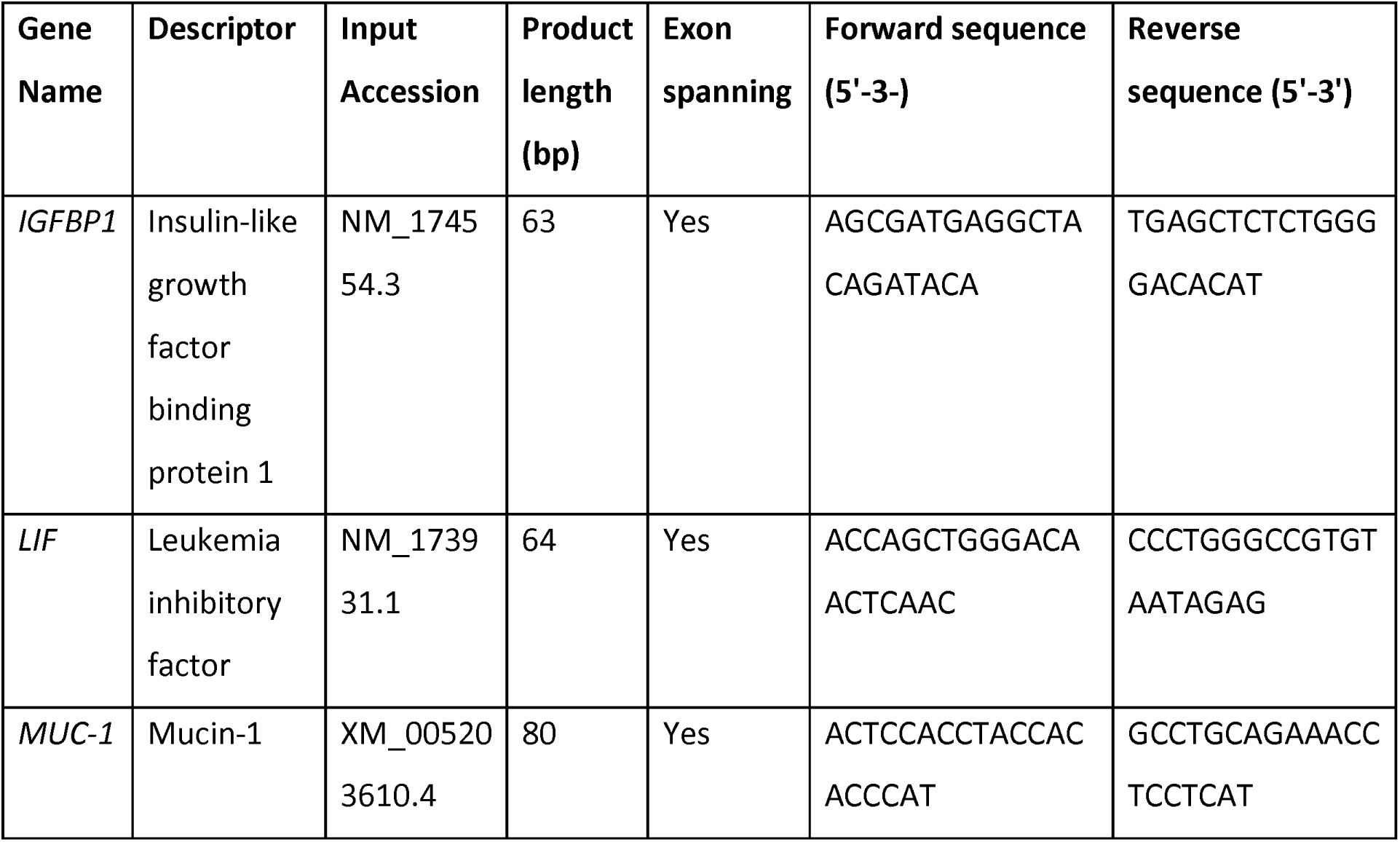
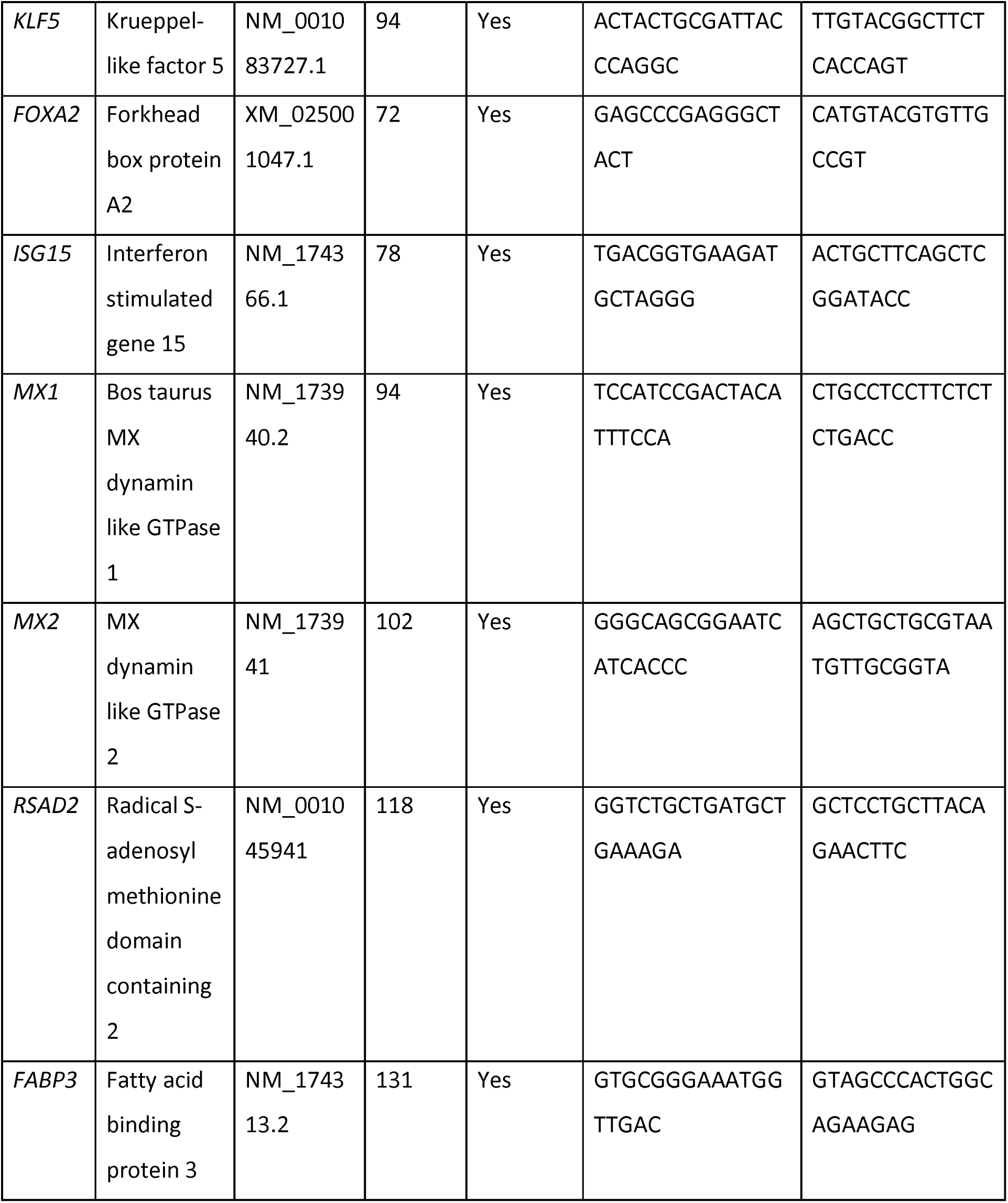
Table of primer sequences for gland markers for initial optimisations and classical and non-classical ISGs. Primers were designed using Primer BLAST for transcripts of known glandular cell origin or classical/non-classical ISGs with a maximum product length of 200bp and exon-exon spanning requirements.

### Culturing of bovine endometrial organoids

Following optimisation steps above, digestive solution 1 (50 ml HBSS, 25 mg Collagenase II, 500 μl 100X Trypsin solution and 125 μL 4% DNAse) was used to fragment tissue for 1 hr, which was then passed through a 100 μM filter, followed by 40 µM filter. The 100 μM filter was discarded, whilst the 40 μM filter was inverted and rinsed with 5 ml of stop solution into a falcon tube. This was centrifuged at 500 x g for 5 min, the supernatant discarded, and the remaining pellet resuspended in 1 ml of Advanced DMEM/F12 medium. This was transferred to an Eppendorf tube and another 5 min centrifugation step (500 x g) was performed. The size of the pellet was estimated and resuspended in a volume of Advanced DMEM/F12 medium relative to pellet size - for a ‘just visible’ pellet approximately 250 μl was used and for a larger, readily visible pellet up to approximately 500 μl was used.

Glandular organoids were then cultured using the same method as previously described^21^. Keeping the Cultrex (Cultrex Reduced Growth Factor Basement Membrane Extract, Type 2, R&D Systems) on ice, 12 μl of resuspended cells was taken for each well and added to 48 μl of Cultrex. After gentle pipetting up and down to mix, a 60 μl drop was placed in the centre of a well of a 24 well plate. The 24 well plate was placed in the incubator for 30 min to set and afterwards each drop was covered with 500 μl of expansion media. The expansion media^21^ is composed of Advanced DMEM/F12 (Life Technologies), N2 supplement (Life Technologies), B27 supplement minus vitamin A (Life Technologies), Primocin 100 μg/ml (Invitrogen), N-Acetyl-L-cysteine 1.25 mM (Sigma), L-glutamine 2 mM (Life Technologies), recombinant human EGF (Peprotech) 50 ng/ml, recombinant human Noggin 100 ng/ml (Peprotech), recombinant human R-Spondin-1 400 ng/ml (Peprotech), recombinant human FGF-10 (Peprotech) 100 ng/ml, recombinant human HGF (Peprotech) 50ng/ml, nicotinamide 10nM (Sigma), and ALK-4, −5, −7 inhibitor (System Biosciences). Full details are provided in Table 2. The medium was replaced every 2-3 days.

**Table 2.** Expansion media components, final concentrations, product numbers and suppliers used for culturing of bovine endometrial organoids. ^21^

| Component | Final Concentration | Product Number | Supplier |
| --- | --- | --- | --- |
| Advanced DMEM/F12 | 1X | 12634010 | Life Technologies |
| B27 | 1X | 12587010 | Life Technologies |
| N2 | 1X | 17502048 | Life Technologies |
| L-glutamine | 2 mM | 25030-024 | Life Technologies |
| N-Acetyl-L-cysteine | 1.25 mM | A9165-5G | Sigma |
| Nicotinamide | 10 nM | N0636 | Sigma |
| Primocin | 100 µg/ml | Ant-pm-1 | Invivogen |
| Stemmacs | 500 nM | ZRD-A8-02 | System Biosciences |
| Recombinant human Rspodin-1 | 400 ng/ml | 120-38 | Peprtech |
| Recombinant human Noggin | 100 ng/ml | 120-10c | Peprtech |
| Recombinant human FGF-10 | 100 ng/ml | 100-26 | Peprtech |
| Recombinant human EGF | 50 ng/ml | AF-100-15 | Peprtech |
| Recombinant human HGF | 50 ng/ml | 100-39 | Peprtech |

### Passaging of organoids

We also sought to determine for how long these organoids could be passaged. To test this, passages were carried out approximately every 10 days (or more at later passages where organoids were slower to expand), once the majority of organoids in the droplet had reached approximately >300 µm. If organoids were allowed to continue beyond this size they would begin to collapse and become dark in the centre. For passaging each well of the culture dish contained 500 μl of expansion media and one droplet consisting of the organoids in cultrex. These droplets were broken up in the well without removing culture media by manual pipetting 4-5 times. Two well’s worth of organoids i.e. 2 technical replicates were transferred to a single Eppendorf and centrifuged at 2400 x g for 6 min. The supernatant was removed and 150 μl advanced DMEM/F12 added, followed by manual pipetting 300 times to break up the organoids and residual Cultrex. One ml of advanced DMEM/F12 was added and tube centrifuged at 600 x g for 6 min before aspiration of supernatant. Another 150 μl advanced DMEM/F12 was added and subjected to manual pipetting a further 100 times before adding an additional 1 ml advanced DMEM/F12 and centrifuging at 1200 x g for 6 min. Following this centrifugation step, the majority of the supernatant was aspirated. The remaining media and organoid mixture was resuspended in multiples of 48 μl of ice-cold Cultrex depending on the number of wells to be seeded in a given passage. Organoids were usually split in a 1:3 ratio. Sixty μl drops were dispensed into the centre of wells of a 24-well plate and this was incubated at 38.5 °C for 30 min to set, followed by addition of 500 μl expansion media.

### Characterisation of organoids via qRT-PCR and Immunoflouresence

In order to characterise the organoids, we examined expression of previously selected gland marker transcripts by qRT-PCR over time. Organoids (n=3) were collected at regular time points by removing expansion media and adding 500 µl fridge cold PBS to soften Cultrex. Contents of the well were pipetted up and down manually 4 times before transfer to an eppendorf for centrifugation at 600 g for 6 mins. Supernatant was then aspirated, and pellet was resuspended in PBS to wash followed by a repeat centrifugation. The supernatant was aspirated again and pellets snap frozen in liquid nitrogen for subsequent RNA extraction using Qiagen MiRNeasy Micro Kit following the manufacturers protocol. Reverse transcription and qRT-PCR were carried out as previously described above. At each time point, organoids were also imaged by light microscopy to assess morphology.

Organoids for immunofluorescence analysis were plated on 13 mm round coverslips within the wells. On collection of these organoids, expansion medium was removed, droplets overlayed with 20 µl Cultrex and placed in the incubator for 20 min to set. Five hundred µl of 4% formaldehyde was then added to the well for 30 min at room temperature. This was then removed and replaced with PBS and stored at 4°C until imaging. Fixative was washed away with PBS before samples were quenched in 50mM ammonium chloride and permeablised in 0.1% triton X-100 for 5 minutes. Samples were then incubated with rabbit anti-E-cadherin (Abcam) or mouse anti-acetylated alpha tubulin antibody (Abcam) diluted 1/100 in PBS overnight at 4°C. Samples were subsequently washed with PBS and incubated for 1 hr at room temperature with Alexa-488-conjugated anti-mouse/rabbit secondary antibodies (Life Technologies) diluted 1/200 in PBS containing 1µg/ml DAPI and 1/50 dilution of Alexa-568-conjugated phalloidin. After washes, stained samples were mounted in mowiol on glass slides and imaged using an apotome-equipped Zeiss Axiophot inverted fluorescence microscope and Zeiss Zen software.

### Treatment of organoids with signals important for early pregnancy

Following isolation of organoids as described above, organoids (derived from n=5 different animals) at passage 0, 8 days post isolation were treated with the following: 1) Control (no treatment), 2) Ethanol Vehicle (70% EtOH), 3) P4 (10 µg/ml progesterone), 4) PBS Vehicle (PBS), or 5) IFNT (1000 ng/ml recombinant ovine IFNT), for 24 hr. After 24 hr, the expansion media was removed and 500 µl of cell recovery solution added. Plates were placed on ice for 1 hr. Well contents were transferred to Eppendorfs, pooling 2 identical wells to ensure sufficient RNA yield. Centrifugation was carried out for 5 min at 500 x g and supernatant discarded. Cells were resuspended in 500 µl of PBS by manual pipetting. An additional centrifuge at 500 x g for 5 min was carried out before supernatant was removed and pellets snap frozen prior to RNA extraction.

### RNA sequencing and data processing

RNA was extracted as described above and sequencing carried out by Novogene. RNA integrity and concentration was determined using the Bioanalyzer 2100 (Agilent Technologies USA) and Nanodrop (Thermo Fisher Scientific, USA). Messenger RNA was purified from total RNA using poly-T oligo-attached magnetic beads. After fragmentation, the first strand cDNA was synthesized using random hexamer primers followed by the second strand cDNA synthesis. The library was prepared using Novogene NGS RNA Library Prep Set (PT042) following end repair, A-tailing, adapter ligation, size selection, amplification, and purification. The library was checked with Qubit and real-time PCR for quantification and bioanalyzer for size distribution detection. Quantified libraries were pooled and sequenced (6GB per sample) on Illumina Novaseq6000 platforms, according to effective library concentration and data amount and using the PE150 sequencing strategy.

Paired-end sequencing quality control was assessed through FastQC (www.bioinformatics.babraham.ac.uk/projects/fastqc) and multiQC^25^. Both adapters and low quality bases (QV < 20) were trimmed from reads’ extremities using Trimmomatic^26^ with a minimum read length of 30 bp. An average of 23.2 million reads per sample were kept post-trimming for downstream analyses. All libraries were mapped against the bovine ARS-UCD1.2 release-109 reference genome (retrieved from https://ftp.ensembl.org/pub/release-109/fasta/bos_taurus/dna/Bos_taurus.ARS-UCD1.2.dna.toplevel.fa.gz) through STAR aligner^27^ with default parameters. STAR-generated sorted BAM output files were used for assigning read counts to gene features with featureCounts^28^ with the following parameters: -p -B -C -M -O --fraction. We relied on Bos_taurus.ARS-UCD1.2.109.gtf annotation file downloaded from the same ensembl ftp address above.

### Differential expression analysis

Differential expression analysis was performed using DESeq2 R package^29^ on data from 25 samples with the following treatment groups (n = 5): (i) Control (Ctrl); (ii) IFNT; (iii) PBS; (iv) P4; (v) Vehicle Control (VC). For visualisation, size factors were estimated from the count data and the Relative Log Expression (RLE) normalisation was used to obtain regularised log transformed values. These normalised values were then used for principal component analysis with plotPCA function in DESeq2 R package. Wald test was used on genes that passed an independent filtering step with the following pairwise comparison groups: IFNT vs PBS; P4 vs VC. Resulting P values were adjusted for multiple testing using the Benjamini-Hochberg procedure. Genes with absolute fold change > 2 and adjusted p-value < 0.05 were considered differentially expressed. Results were visualized in a volcano plot using ggplot R package. All tools described in this paragraph were run under the R environment version 4.4.0.

### Functional enrichment analysis using DAVID and Enrichment Map

Selected genes from differential expression analysis were used for gene-set functional enrichment analysis with DAVID tool^30^, setting species to Bos taurus. For each comparison pair, total number of genes and separately up- and downregulated genes were analysed. EASE score (modified Fisher Exact p-value of enrichment) was set to 0.05. Functional enrichment network was built based on DAVID output charts of gene-set enrichment for each comparison pair using EnrichmentMap app (v. 3.5.0)^31^ in Cytoscape software (v.3.10.3)^32,33^ with edge cutoff set to 0.375 (merged Jaccard and Overlap coefficients). Autoannotate App (v.1.5.1) with MLC algorithm based on similarity coefficient was used to create annotated groups.

### In silico protein-protein interaction analysis using STRING

In silico protein-protein interaction analysis of selected genes was performed on the basis of the STRING database for Bos taurus^34,35^. Interaction networks were built based on the list of selected genes from each comparison pair using stringApp (v.2.1.1)^36^ in Cytoscape with confidence cutoff score set to 0.4. Functional enrichment of formed clusters was performed using *Bos taurus* genome as a background, enriched terms were analysed with varying redundancy cutoff settings.

### Expression of a conserved panel of microRNAs in bovine endometrial organoids in response to P4 and IFNT treatment

To test if these organoids can be used to assess key non-coding RNA molecules important for implantation, we investigated expression of a conserved set of microRNAs previously shown to be important for implantation^37^. MicroRNA expression in endometrial organoids treated with P4 and IFNT (as described above) was investigated using RNA from animals 1, 3 and 4. MiRCURY LNA RT Kit (Qiagen) was used - following manufacturers’ instructions - to reverse transcribe cDNA for qRT-PCR. Each reaction contained 10 ng/µl RNA. Custom miRCURY LNA miRNA PCR Panels (Qiagen, Cat #339330) were used which are preloaded with primers for conserved eutherian mammal specific miRNAs of interest^21,37^ and samples carried out in technical duplicate. Plates were run on the LightCycler 96 (Roche) PCR machine with the following cycle: 95 °C for 2 min; 40 cycles of 95 °C for 10 sec and 56 °C for 1 min; followed by a melt curve analysis from 60-95 °C. Cq values were obtained via LightCycler 96 software and transferred to Microsoft Excel for processing. ΔCt values were normalised by subtracting the 5S normaliser from each value. The transformation 2^-ΔCt^ was calculated for each value and transferred to GraphPad Prism for graphical representation. A one-way ANOVA with multiple comparisons was performed on the values to determine statistical significance.

### Culturing of organoids in a microfluidics device

Microfluidic devices were fabricated by soft lithography in polydimethylsiloxane (PDMS, Sylgard® 184, Dow Corning, MI, USA). Liquid PDMS (10:1) was casted and cured on SU8 2100 moulds (Microchem, MA, USA). SU8 2100 photoresist was spun on a silicon wafer at 2000 rpm for 30 sec to obtain a 150 μm thick layer. Wafer covered with SU8 was exposed to UV light (wavelength= 375nm, energy=250mJ/cm^2^) through a film photomask designed using computer-aided design (CAD) software (AutoCAD). Once cured, the PDMS layer was peeled off from the SU8 mould and inlet and outlet ports were opened using a 1.5mm round puncher (Integra Mylex, NJ, USA). The device was then assembled by treating a glass slide (2.54 x 5.08 cm) and the PDMS layer by Oxygen Plasma (600mTorr, 100W, 30s). 500 μL Pyrex cloning cylinders (Fisher Scientific, Pittsburg, PA, US), used as reservoirs for cell media, were bonded to the port regions using liquid PDMS. Prior to use, devices were sterilized by UV exposure (wavelength= 254nm for 25 min). Briefly, the device consisted of a rhomboidal shaped chamber (area: 48.85 mm^2^) with 60° angled inlet and outlet microchannels. The design includes round PDMS pillars (diameter 100 μm) to provide structural support for the culturing chamber. For the culture of the organoids, the chamber was filled with Polyvinylpyrrolidone (PvP), subsequently the organoids were loaded into the device and then left for 3 days in the incubator to set. After this time, they were treated with IFNT as previously described.

## RESULTS

### Optimisation of Cell Isolation Process

In order to select the optimal protocol for isolation of bovine endometrial epithelial gland cells, two time points (30 min and 1 hr) and three digestion compositions were used, as well as collection of different fractions of digestion (Figure 1C) to be analysed. Isolation and culture of glands resulted in similar organoid gland morphology to that described in other species^21^ (Figure 1B). Gland marker gene expression *IGFBP1, LIF, MUC-1, FOXA2* and *KLF5* were measured by qRT-PCR (Figure 2A-E). There was consistently lower gland marker expression for all genes in cells obtained from digestion composition 3, as well as absence of each gland marker from at least 1 sample. Cells isolated using digestion 1 and 2 displayed similar, and higher (compared to digestion 3) gland marker expression values for each gene, which resulted in selection of Digestion 1 due to fewer components and it being consistent with the ability to isolate stromal cells from the same individual. Cells isolated from the 70 μM strainer show reduced gland marker expression for all animals at both 30 min and 1 hr timepoints compared to cells isolated from the 40 μM strainer. For this reason, cells collected from inverting the 40 μM strainer were selected for use. All gland markers exhibit higher expression following 1 hr incubation compared to 30 min. Overall, conditions selected were: 1 hr incubation with Digestion mix 1, using cells taken from the 40 μM cell strainer (Figure 1D).

**Figure 2:**
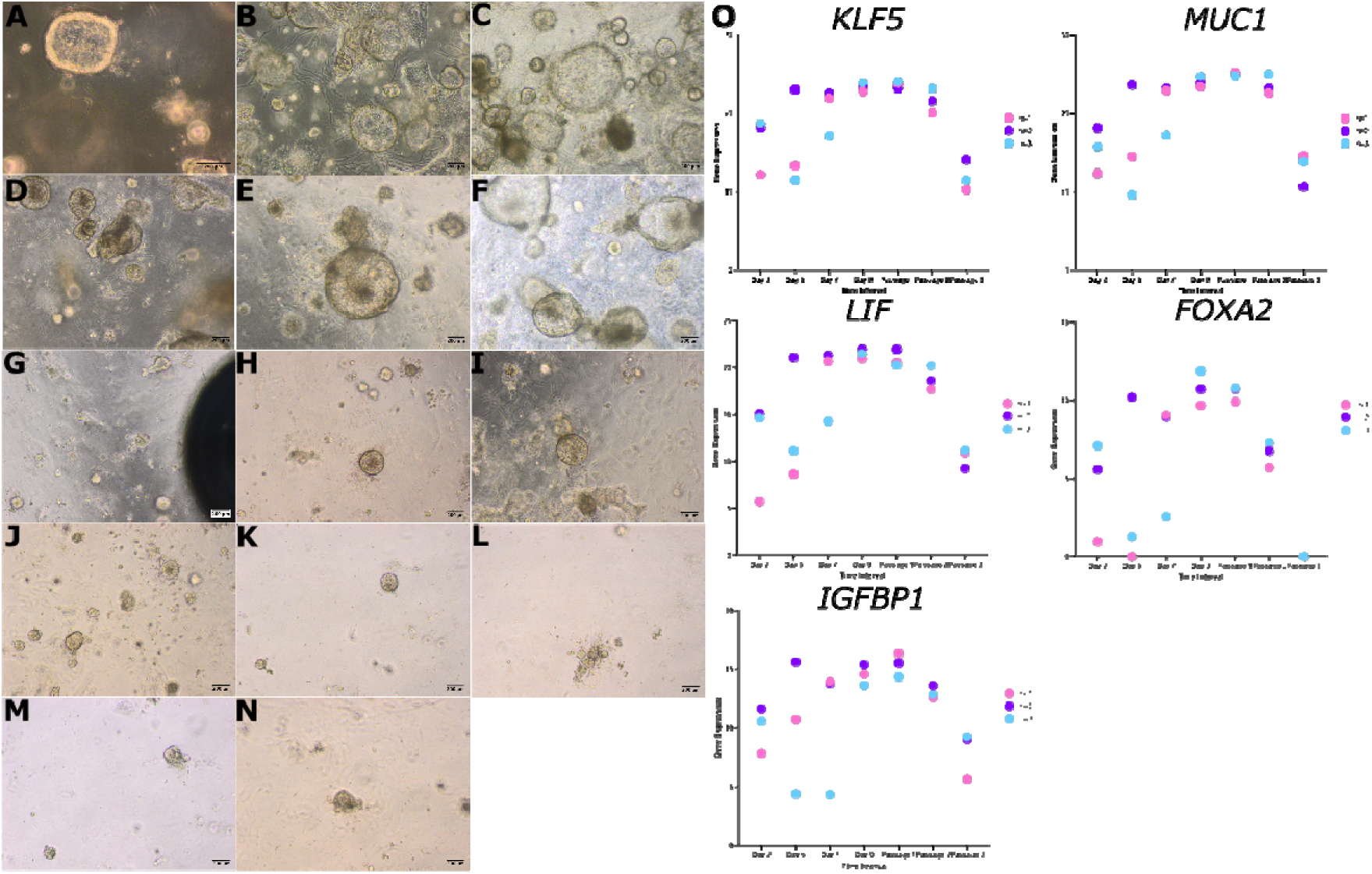
Representative light microscopy images of bovine endometrial organoids (n=2) at different time points of growth. **(A)** Day 5, **(B)** Day 7, **(C)** Day 9, **(D)** Passage 1 + 5 days, **(E)** Passage 1 + 7 days **(F)** Passage 1 + 10 days. **(G)** Passage 2 + 2 days, **(H)** Passage 2 + 7 days, **(I)** Passage 2 + 10 days, **(J)** Passage 2 + 15 days. **(K)** Passage 3 + 3 days, **(L)** Passage 3 + 7 days, **(M)** Passage 3 + 10 days, **(N)** Passage 3 + 14 days**. (O)** Raw gene expression (raw CT value subtracted from 45 - maximum CT value) of gland markers KLF5, MUC1, LIF, FOXA2 and IGFBP1 in bovine endometrial organoids (n=3) at the following points of growth post isolation: day 2, day 7, day 9, passage 1, passage 2 and passage 3.

### Response to passaging over time

Bovine endometrial organoids (n=3) were cultured with imaging at regular time points to evaluate growth and passaging outcomes. Example images of organoids from one animal are shown in Figure 2. Images for other animals (n=3) can be found in supplementary materials (Supplementary Figure 1 and 2). Similar trends can be identified in all animals. All three exhibit growth during passage 0 and 1, with animal 2 developing larger organoid structures than animal 1. Growth rate reduces at passage 2 and organoids do not expand to the size they previously reached. Growth is poor at passage 3 where there is little to no growth in any animal at any stage.

Gland marker expression was measured by qRT-PCR over three passages to characterise gene expression in the organoids. Raw Cq values were used to test gland marker expression as a preliminary method of detection of expression. All 5 genes: *KLF5, MUC1, LIF, FOXA2* and *IGFBP1* were expressed in all animals (n=3) but not necessarily at all time points (Figure 2O). *FOXA2* was not expressed at passage 3 in any animal and in fact this gene exhibits varied expression between animals at earlier stages. At very early stages (day 2, 5 and 7) for all genes, expression appears to vary dramatically between the animals. Upon reaching day 9 and during passage 1, gene expression of these 5 markers becomes similar. Expression of all 5 genes is reduced during passage 3.

### Localisation of markers of organoids

Fluorescence microscopy analysis of bovine endometrial organoids the confirmed apicobasal polarity of the cells, with actin staining strongly present at the apical domain facing the inner lumen of the organoids and anti-E-cadherin staining localised to basolateral cell membranes (Figure 3A). Anti-acetylated alpha-tubulin antibody was used to identify ciliated cells, however the antibody staining intracellular tubulin fibres only with no apically located cilia visible (Figure 3B).

**Figure 3.**
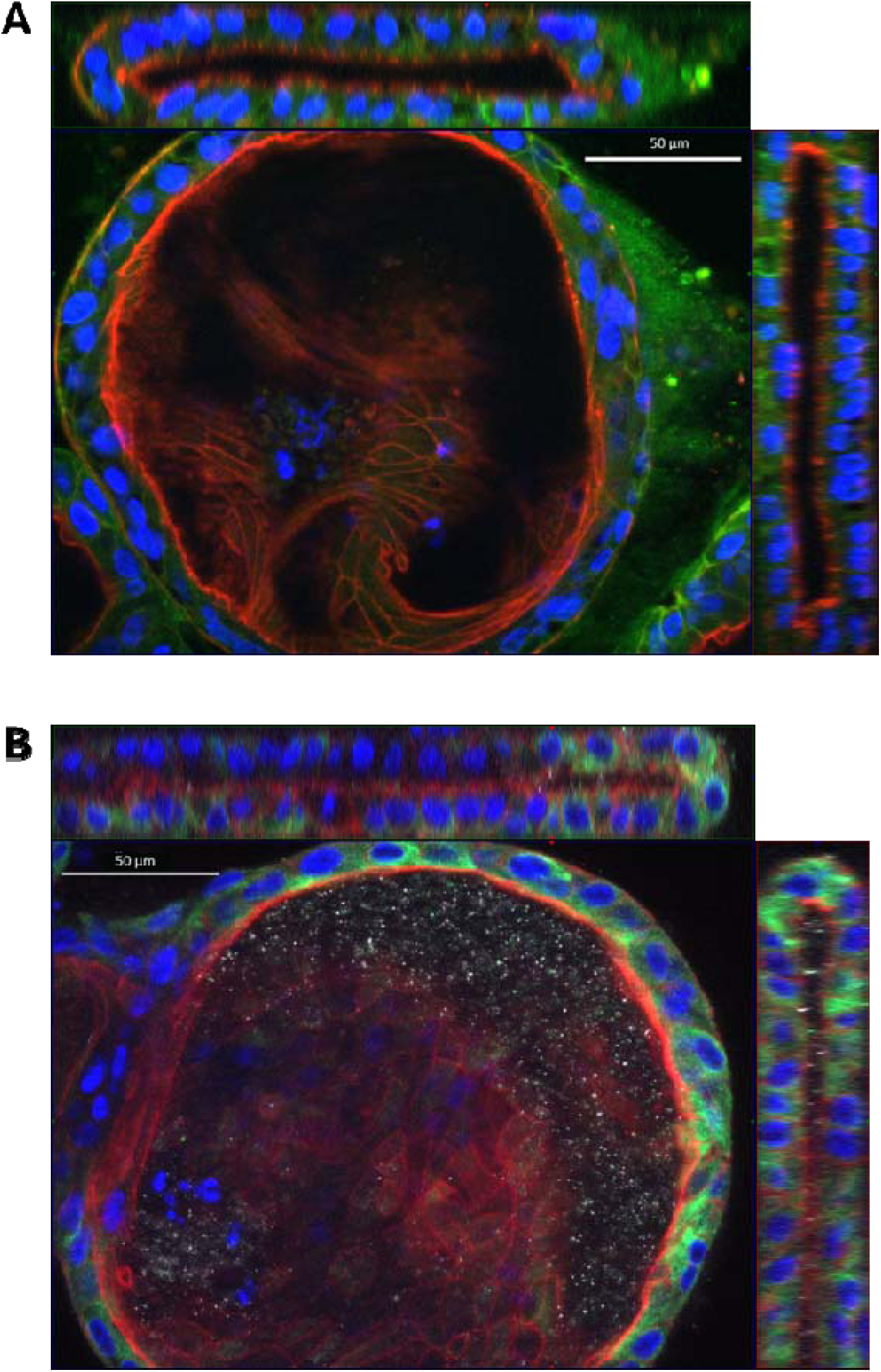
Epithelial morphology of bovine endometrial organoids. 3D fluorescence microscopy of **(A)** anti-E-cadherin (green), or **(B)** anti-acetylated alpha-tubulin with phalloidin (red) and DAPI (blue) stained bovine endometrial organoid. Main panel shows x-y cross section through the centre of the organoid and upper and rightmost panels show x-z and y-z cross sections, respectively.

### Transcriptional response of bovine organoids to Progesterone

Principal component analysis (PCA) demonstrated that biological replicate was responsible for the largest variance observed in the data (Figure 4A), however a clear separation of P4 treated organoids and vehicle or control treatments is evident. Again, control samples cluster with corresponding vehicles suggesting the vehicle has minimal effect on detected differential expression. Following 24h treatment with P4, there was a total of 169 significantly upregulated genes, with 102 significantly downregulated transcripts (Figure 4B; Supplementary Table 1). Functional enrichment analyses of these genes were carried out and protein partner interaction networks were generated. For transcripts altered by P4, there is clear clustering of genes involved in cell migration, extracellular matrix and transmembrane transport (Figure 4). Full network is shown in Supplementary Figure 3, and Supplementary File 1 provides an interactive view of the networks and data tables.

**Figure 4.**
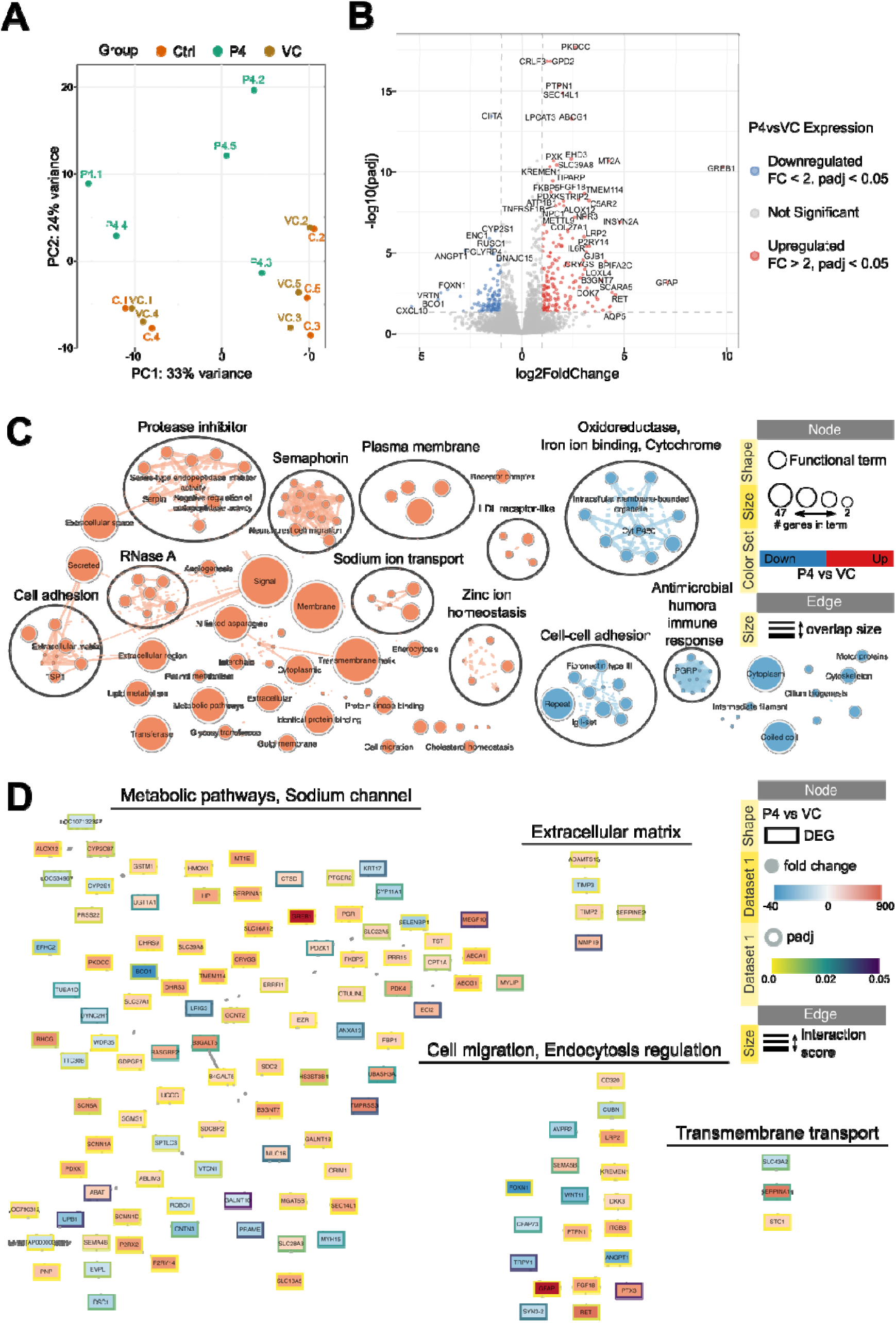
Effect of progesterone treatment on bovine endometrial organoids. **(A)** Principal component analysis plot representing variation in samples from control and treatment groups. Ctrl – Control; VC – vehicle control. **(B)** Volcano plot showing differentially expressed genes in P4 vs VC samples (Wald test, padj <0.05, |FC| > 2). Padj – adjusted p-value; FC – fold change. **(C)** Functional terms and pathways enriched in samples from treatment or control groups. Enrichment map displays enriched gene-sets based on differentially expressed genes in P4 vs VC organoids. **(D)** STRING network of predicted protein interactions for genes responding to P4. The Markov Cluster Algorithm (MCL) was used for clustering based on interaction score, with granularity set to 2 and edge weight cutoff set to 0.4. Protein interaction clusters were analysed for functional enrichment within StringApp; only selected clusters are shown. Supplementary File 1 contains the web session of both networks for interactive view.

### Transcriptional response of bovine organoids to IFNT

A PCA plot (Figure 5A) demonstrates that the largest source of variance was the transcriptional response to IFNT, with IFNT treated organoids clustering to the right of the plot and vehicles and their respective controls clustering to the left. Treatment with IFNT resulted in a total of 414 upregulated and 119 downregulated transcripts (Figure 5B, Supplementary Table 2). To understand the functional relevance of differentially expressed genes we undertook functional enrichment analyses and generated predicted protein-protein interaction networks. Figure 5C shows enriched functions, pathways and cellular components for up (red) and down (blue) regulated genes. Selected clusters of predicted protein-protein interactions are displayed in Figure 5D; Supplementary Figure 4 shows full network. For those transcripts altered by treatment with IFNT, the largest cluster is associated with defence to virus and interferon signalling, with others relating to retinol metabolism, serine protease inhibitor and ion transport. Supplementary File 2 provides an interactive view of the networks and data tables.

**Figure 5.**
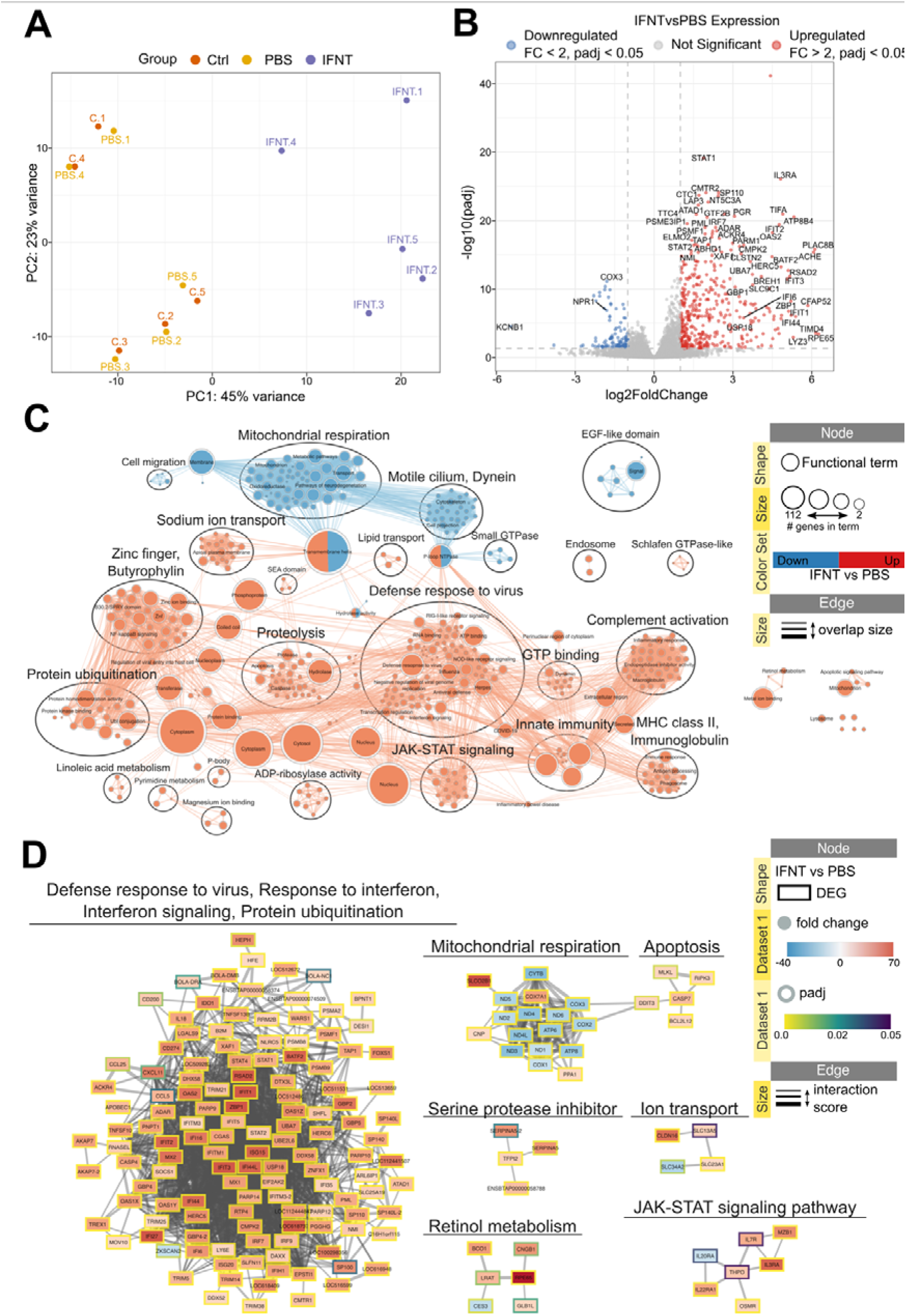
Effect of IFNT treatment on bovine endometrial organoids. **(A)** Principal component analysis plot representing variation in samples from control and treatment groups. Ctrl – Control; VC – vehicle control. **(B)** Volcano plot showing differentially expressed genes in IFNT vs PBS samples (Wald test, padj <0.05, |FC| > 2). Padj – adjusted p-value; FC – fold change. **(C)** Functional terms and pathways enriched in samples from treatment or control groups. Enrichment map displays enriched gene-sets based on differentially expressed genes in P4 vs VC organoids. **(D)** STRING network of predicted protein interactions for genes responding to IFNT. The Markov Cluster Algorithm (MCL) was used for clustering based on interaction score, with granularity set to 2 and edge weight cutoff set to 0.4. Protein interaction clusters were analysed for functional enrichment within StringApp; only selected clusters are shown. Supplementary File 2 contains the web session of both networks for interactive view.

### Comparison of IFNT and P4 transcriptional response in bovine organoids

Of the transcripts altered by IFNT or P4 in the organoids, 30 of them were shared between both treatments (Figure 6A). Twelve of these genes are altered in the same way (e.g. increased expression in response to P4 or IFNT treatment) in response to both treatments, and 18 in the opposite way (e.g. increased expression following P4 treatment and decreased expression following IFNT treatment). There were several enriched functional processes and pathways which involve a combination of genes altered by IFNT and those altered by P4. For example, the oxidoreductase cluster in the enrichment map involves genes from both IFNT (purple) and P4 (green) (Figure 6B). In predicted protein-protein interaction network, sodium channel cluster involves Sodium Channel Epithelial 1 Subunit Alpha (SCNN1A), which is altered by both IFNT and P4, along with 4 genes altered by P4 and 5 genes altered by IFNT (Figure 6C, Supplementary Figure 5). Collagen formation, and extracellular matrix is another cluster which exhibits multiple genes from each list functioning in the same cluster. Supplementary File 3 provides an interactive view of the networks and data tables.

**Figure 6.**
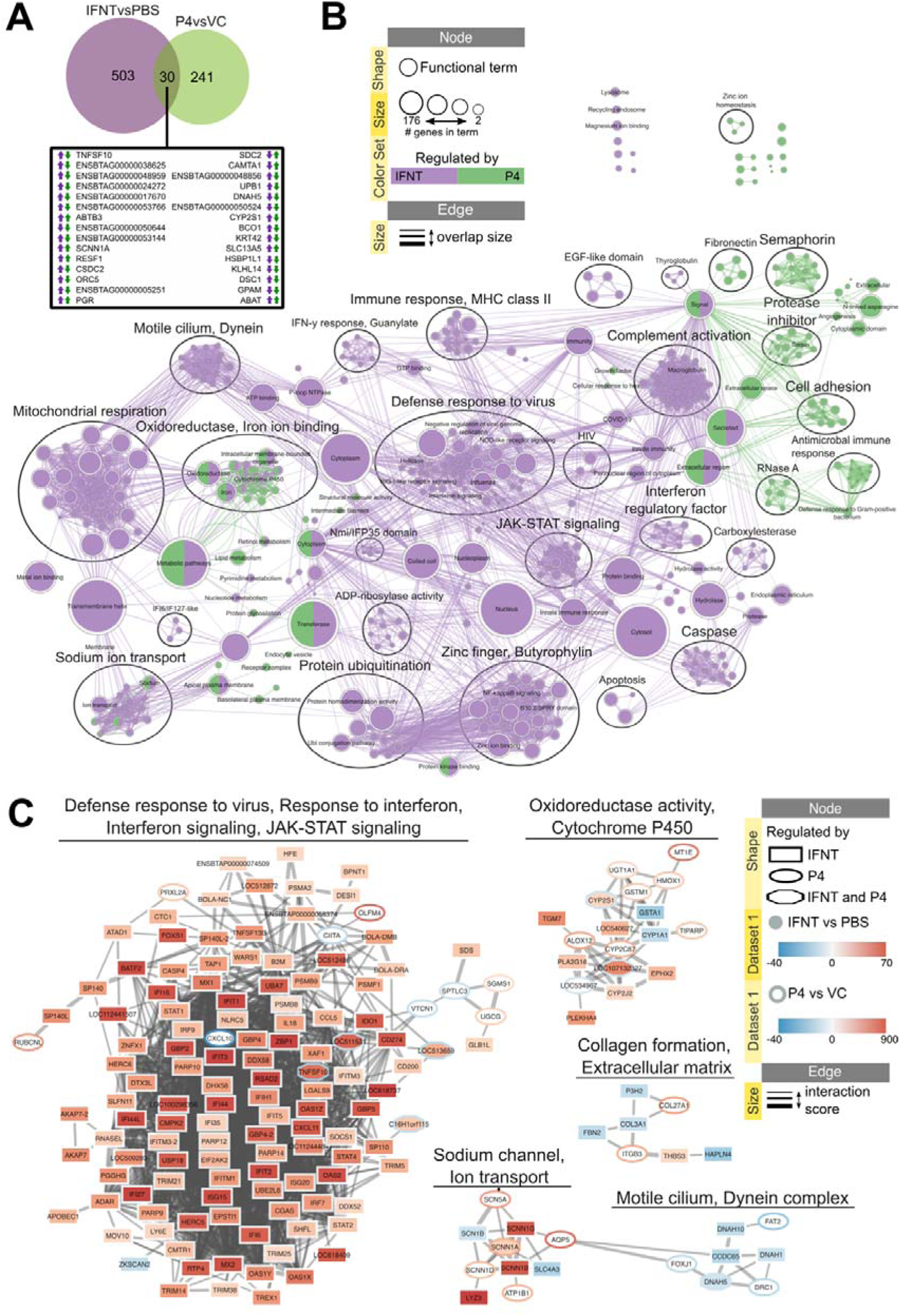
Separate and shared effects of P4 and IFNT treatment on bovine endometrial organoids. **(A)** Venn diagram depicting shared differentially expressed genes from P4 vs VC and IFNT vs PBS comparisons. **(B)** Functional terms and pathways enriched in organoids after P4 or IFNT treatments. Enrichment map displays enriched gene-sets based on differentially expressed genes in P4vsVC and IFNTvsPBS organoids. **(C)** STRING network of predicted protein interactions for genes responding to P4 or IFNT. The Markov Cluster Algorithm (MCL) was used for clustering based on interaction score, with granularity set to 2 and edge weight cutoff set to 0.4. Protein interaction clusters were analysed for functional enrichment within StringApp; only selected clusters are shown. SupplementaryFfile 3 contains the web session of both networks for interactive view.

### Potential applications of bovine endometrial organoids

To investigate the future applications of this *in vitro* model, we examined expression of microRNAs conserved across placental mammals ^37^. We detected expression of all 20 of these miRNAs in our endometrial organoids. Following this, we measured expression of the miRNAs in organoids treated with IFNT and P4. Treatment with 10 μg/ml P4 resulted in significantly (p<0.05) reduced expression of both miR-378a-3p and miR-708-5p compared to control (Supplementary Figure 6). However, there was no significantly differential expression compared with the vehicle control (ethanol) suggesting that effects could be due to vehicle rather than P4. There was also no differential expression of the miRNAs in response to IFNT treatment (Supplementary Figure 7). Nonetheless, this data demonstrates that conserved microRNAs are expressed in the organoids.

As future approaches will seek to culture these organoids in microfluidic devices, we cultured and treated organoids in these devices (Figure 7B), without flow. Firstly, it was possible to culture organoids in a microfluidic device: fragmentation and morphological changes were not observed, excluding detrimental effects of the pressure during loading and growth. However, treatment within the device with IFNT in static conditions, resulted in increased expression of *MX1* (P<0.001), *ISG15* and *RSAD2* (P<0.01), but there was no effect of IFNT on expression of *MX2* (P>0.05) compared to control. For the non-classical *ISGs IGFBP1*, and *FABP3* there were no effects of IFNT on expression of these transcripts (P>0.05) compared to non-stimulated groups (Figure 7C).

**Figure 7.**
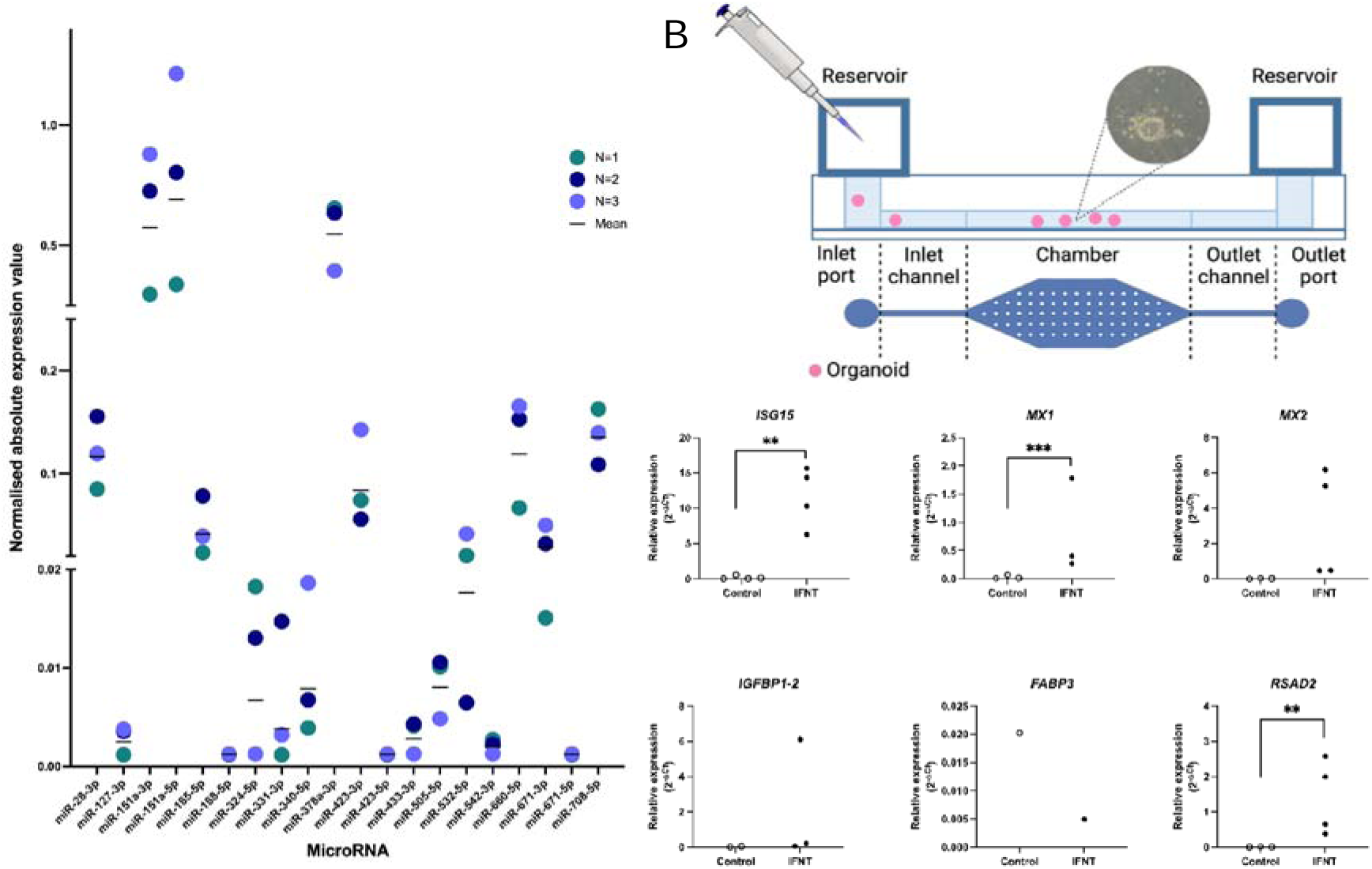
Potential applications for bovine endometrial organoids as an in vitro model. **(A)** Expression of selected microRNAs of interest in bovine endometrial organoids. Organoids (n=3) were cultured for 9 days with no treatment and microRNA expression investigated by quantitative real-time PCR, with values normalised to 5s. **(B)** Images comparing morphology of bovine endometrial organoids cultured in traditional static well conditions (top) and static microfluidic devices (bottom). **(C)** Quantitative real-time PCR analysis of selected canonical (*ISG15, MX1, MX2, RSAD2)* and non-classical ISGs (*IGFBP1, FABP3*) in organoids treated with IFNT (recombinant ovine IFNT 100 ng/µl) for 24 hrs in a microfluidic device under static conditions. Black circles represent gene expression in organoids following treatment with IFNT, white circles represent gene expression in organoids following PBS Control treatment. Significant differences in gene expression are marked with an asterix (*) when p<0.05 or two asterix (**) when p<0.01. Analysis was carried out using paired t-test.

## DISCUSSION

Here, we have produced novel 3D glandular epithelial endometrial organoids from bovine endometrium which respond to key pregnancy molecules P4 and IFNT. Glandular endometrial organoids are a great model for studying endometrial function and early pregnancy events such as implantation, as uterine gland secretions are thought to be vital for implantation and fertility^38^. We also demonstrate that these organoids express gland marker genes, conserved microRNAs, and can be cultured in microfluidic devices - vastly increasing the future research opportunities for this model. Moreover, these respond to P4 and IFNT and demonstrate a deep glandular phenotype. Collectively these data show a novel way to study specific aspects of endometrial function in cattle.

The full extent of the essential interactions between conceptus and uterus required to support establishment of a successful pregnancy have yet to be fully elucidated. Organoids have been successful in different human systems including endometrium^22^ and adding this form of complexity to the bovine endometrial system could help tailor a more physiological approach to investigate interactions *in vitro*. In our model, variation can be observed in growth rate and organoid size over time between animals. To some degree this highlights animal to animal variation which is to be expected, and has been previously shown in other in vitro bovine models^15,39^. However, this could also be attributed to plating density, as cell pellet volume was estimated when resuspending to plate the cells. Furthermore, for the purpose of this experiment, each animal’s organoids were passaged on the same day irrespective of the organoid size, which may not have been at an optimal stage for some animals. It is possible that for some animals, the organoids never reached the appropriate size or cell density and so remained consistently smaller throughout the time course. A drawback of this model is the limited longevity of the organoids, with growth halting at passage 3. This result suggests that these organoids should be used for experiments during earlier passages for optimal results. However, potential changes to the modify expansion media for species specificity may help with further passaging given this media was optimised for human endometrial organoid culture^21^. Future work to improve this model would be to analyse and optimise media composition which may improve growth and allow the organoids to persist in culture further than passage 3.

Candidate gland marker expression profile was quantified in the organoids to show presence or absence of gland markers (Figure 2O). At many stages of growth, the expression appears very similar between animals despite there being no normalization of the data. Large differences between gland marker expression at early stages could be due to variation in organoid growth rate and differences in plating densities.

### Response to cues

We have shown that these organoids respond to some key cues important for early pregnancy. The organoids are P4 responsive, with many genes that are differentially expressed following exposure linked to and enriched in important biological processes during early pregnancy events. The gene exhibiting highest upregulation following P4 treatment is *GREB1* (growth regulating estrogen receptor binding 1). Fascinatingly, this gene in humans is characterised as a P4 regulated gene^40^; however, it is found to be necessary for stromal cell decidualization, which does not occur in non-invasive implantation as in bovine^41^. In other systems, *GREB1* is linked to oestrogen upregulation^42^ and hormone induced growth of breast cancer^43^ and prostate cancer^44^. Therefore, it is possible that P4 upregulation of *GREB1* in bovine organoids is important for growth and proliferation of the endometrium. Janus kinase 3 (*JAK3*) is another upregulated gene following P4 treatment, which modulates immune response; and an increased immune response has been observed when *JAK3* is inhibited in the intestine^45^. Correct control of the immune response during early pregnancy to allow tolerance of the developing, genetically different offspring is vital. Considering that P4 is elevated during implantation and pregnancy, in turn upregulating expression of *JAK3*, this may be a mechanism to reduce immune response against the conceptus.

From our data we demonstrate bovine endometrial organoids also respond to the pregnancy recognition signal IFNT, with those transcripts that are differentially expressed implicated in processes important for early pregnancy. Defence to virus is the pathway containing the largest number of differentially expressed genes demonstrating that bovine organoids have a classical ISG response to IFNT. Genes that have classical ISG response in the organoids include upregulation of *MX1, MX2* and *ISG15*. Ion transport is another biological process enriched in genes altered by IFNT treatment. As the ULF is known to contain ions^46^, and the organoids display deep glandular phenotype, this may indicate that conceptus produced IFNT acts upon the deep gland cells to alter composition of the ULF in order to support conceptus development and implantation. STRING interaction networks also highlight retinol metabolism, with differentially expressed genes involved in this process significantly upregulated with high fold changes. Retinol metabolism is important for correct development of the conceptus during early development, and it is likely that this process is involved in interactions between the endometrium and conceptus.

When comparing response of the glandular endometrial organoids to IFNT and P4, 30 transcripts are changed by both treatments. Enrichment maps and STRING networks are interesting as they demonstrate the interaction between the genes altered by each or both treatments, as both of these molecules are present during the peri-implantation period. According to transcriptional response by the organoids, it is clear these molecules have specific and different roles when interacting with these deep glandular phenotype cells of the endometrium, with only a small number changing the same pathways and networks. For example, collagen formation and extracellular matrix is highlighted as a network in which genes from both treatments interact. Extracellular matrix remodelling is a vital process during early stages of pregnancy to support implantation and subsequent placentation, with dynamic changes in extracellular matrix protein abundance throughout^47^. It appears P4 and IFNT contribute to this by acting upon the deep glandular epithelial cells of the endometrium. In addition, the differences in response of these organoids to P4 and IFNT likely reflect the fact this is a deep gland phenotype of the organoids i.e. they retain expression of progesterone receptor and have a classical ISG response compared to e.g. the luminal epithelial cells during pregnancy recognition.

Our microfluidic experiment shows that it is possible to culture bovine endometrial organoids within microfluidic chambers which will allow experiments utilising this model to be carried out under controlled dynamic fluid exchange and tuned shear stress. The allows us to test hypotheses in an environment that better mimics the in vivo scenario. In addition, organoids have also been treated with IFNT within these devices, and the results indicate that this upregulated expression of genes that are increased in deep uterine glands during the pregnancy recognition process in ruminants i.e. it mimics the in vivo response^48^. This increase in expression was found for both static culture and culture within microfluidic devices. This is important as the endometrium responds to the pregnancy recognition signal. In vivo, both classical and non-classical *ISGs* are stimulated in the different cells of the endometrium^49^. We show that these organoids express classical ISGs following IFNT stimulation but not non-classical ISGs. This could be due to duration of exposure as previous in vitro approaches have shown that 24 hr exposure is sufficient to induce classical *ISGs* but non-classical requiring more sustained exposure^50^. Another major benefit of this protocol is for the possibility to isolate luminal epithelium and stromal cells as well as organoids from the same animal. This will facilitate future complex building of *in vitro* models incorporating all three cell types.

Interestingly, many of the miRNAs identified to be conserved across placental mammals^37^ are expressed in these bovine endometrial organoids. Whilst none of the miRNAs exhibited differential expression in response to IFNT or P4, these data would be more conclusive with a larger sample size due to the variation between individuals which is already a known issue. In response to P4 treatment, the vehicle has a large effect on expression of many of the miRNAs. This being 100% ethanol, a different vehicle such as water or DMSO would be beneficial to examine differential expression in response to P4 without the influence of the vehicle.

Future approaches will involve isolation of secretome from the organoids, culture under flow conditions to examine transcriptional response in a more physiologically accurate model, knockout of specific genes via CRISPR/CAS9, as well as optimisation of culture components to be species-specific. Furthermore, incorporation of stromal cells to produce endometrial assembloids which has already been achieved in humans (Rawlings Makwana Taylor et al 2021) would assist in creating an *in vitro* multicellular model of the bovine endometrium for investigating early pregnancy events.

In conclusion, we have characterised a bovine endometrial organoid culture method that results in expression of markers of endometrial glands and has important organoid features. Moreover, we show it responds to the two key molecules responsible for changing the transcriptomic landscape of the bovine endometrium during early pregnancy^51^. Collectively these data show a novel 3D method to investigate events in early pregnancy including implantation, endometrial function and communication between the mother and conceptus.

## ACKNOWLEDGEMENTS

This work was supported by BBSRC grant numbers BB/R017522/1, BB/X007367/1 and AN2944 Interdisciplinary DV Awards to NF. We would like to acknowledge the assistance of University of Leeds LeedsOmics facility in RNASeq data analysis, and Biorender in production of Figure 1C. Data in the process of being uploaded to GEO.

