## Supplementary Figure Legends for "Bovine endometrial organoids: A new tool to study conceptus-maternal interactions in mammals"

Supplementary Figure 1: Light microscopy images of bovine endometrial organoids (n=1) at different time points of growth. **(A)** Day 5, **(B)** Day 7, **(C)** Day 9, **(D)** Passage 1 + 5 days, **(E)** Passage 1 + 7 days **(F)** Passage 1 + 10 days. **(G)** Passage 2 + 2 days, **(H)** Passage 2 + 7 days, **(I)** Passage 2 + 10 days, **(J)** Passage 2 + 15 days. **(K)** Passage 3 + 3 days, **(L)** Passage 3 + 7 days, **(M)** Passage 3 + 10 days, **(N)** Passage 3 + 14 days**.**

Supplementary Figure 2: Light microscopy images of bovine endometrial organoids (n=3) at different time points of growth. **(A)** Day 5, **(B)** Day 7, **(C)** Day 9, **(D)** Passage 1 + 5 days, **(E)** Passage 1 + 7 days **(F)** Passage 1 + 10 days. **(G)** Passage 2 + 2 days, **(H)** Passage 2 + 7 days, **(I)** Passage 2 + 10 days, **(J)** Passage 2 + 15 days. **(K)** Passage 3 + 3 days, **(L)** Passage 3 + 7 days, **(M)** Passage 3 + 10 days, **(N)** Passage 3 + 14 days**.**

Supplementary Figure 3: Effect of P4 treatment on bovine endometrial organoids. STRING network of predicted protein interactions for genes responding to P4. The Markov Cluster Algorithm (MCL) was used for clustering based on interaction score, with granularity set to 2 and edge weight cutoff set to 0.4. Protein interaction clusters were analysed for functional enrichment within StringApp. Supplementary file 1 contains the web session of the network for interactive view.

Supplementary Figure 4: Effect of IFNT treatment on bovine endometrial organoids. STRING network of predicted protein interactions for genes responding to IFNT. The Markov Cluster Algorithm (MCL) was used for clustering based on interaction score, with granularity set to 2 and edge weight cutoff set to 0.4. Protein interaction clusters were analysed for functional enrichment within StringApp. Supplementary file 2 contains the web session of the network for interactive view.

Supplementary Figure 5: Separate and shared effects of P4 and IFNT treatments on bovine endometrial organoids. STRING network of predicted protein interactions for genes responding to P4 or IFNT. The Markov Cluster Algorithm (MCL) was used for clustering based on interaction score, with granularity set to 2 and edge weight cutoff set to 0.4. Protein interaction clusters were analysed for functional enrichment within StringApp. Supplementary file 3 contains the web session of the network for interactive view.

Supplementary Figure 6: Effect of P4 on expression of selected microRNAs of interest in bovine endometrial organoids. Bovine organoids (n=3) treated for 24 hrs with 1) control, 2) vehicle - ethanol (volume equal to that of progesterone treatment added to well) or 3) 10μg/ml progesterone. Expression quantified by qRT-PCR and values normalised to 5s. Differences in expression were determined using an ANOVA with Tukey's multiple comparisons test where statistical significance was met when the adjusted p value <0.05.

Supplementary Figure 7: Effect of IFNT on expression of selected microRNAs of interest in bovine endometrial organoids. Bovine organoids (n=3) treated for 24 hrs with 1) control - PBS (volume equal to that of IFNT treatment added to well) or 2) 100 ng/µl recombinant ovine IFNT. Expression quantified by qRT-PCR and values normalised to 5s. Differences in expression were determined using an ANOVA with Tukey's multiple comparisons test where statistical significance was met when the adjusted p value <0.05.
